# Dynamic respiration-neural coupling in substantia nigra across sleep and anesthesia

**DOI:** 10.1101/2025.06.11.659127

**Authors:** Kolsoum Dehdar, Elliot Neuberg, Bon-Mi Gu

**Affiliations:** Neuroscience Institute, HMH JFK University Medical Center, Edison, NJ 08820, USA; Department of Neurology, Hackensack Meridian School of Medicine, Nutley, NJ 07110, USA

## Abstract

Respiration is increasingly recognized as a coordinator of neural activity across widespread brain regions and behavioral states. Even during sleep, respiration rhythms modulate sleep-related oscillations. While the basal ganglia are known to play roles in both sleep and respiratory regulation, their interaction with respiration rhythms remains poorly understood. Here, we examined respiration-neural couplings in the substantia nigra pars reticulata (SNr), a major output nucleus of the basal ganglia, and the primary motor cortex (M1) across multiple states in male and female mice, including non-rapid eye movement (NREM) sleep, rapid eye movement (REM) sleep, quiet wakefulness, and anesthesia. Simultaneous recordings of local field potentials (LFPs) from M1 and SNr along with diaphragm muscle activities revealed state-dependent, region specific patterns of respiration-neural coupling. Coupling strength in both SNr and M1 was attenuated during NREM sleep compared to REM sleep and quiet wakefulness. However, under ketamine/xylazine anesthesia, coupling was markedly enhanced in the SNr, but not in M1, indicating region-specific sensitivity to arousal and anesthesia state. Notably, respiration-neural coupling was systematically related to delta sub-band power; coupling strength was reduced with increased slow delta (0.5-2 Hz) and decreased fast delta (2.5-4 Hz) powers. In addition, slow delta was associated with SNr-M1 synchronization, suggesting that inter-regional communication during deep sleep may suppress respiration locking. Together, these findings highlight dynamic, state-dependent modulation of respiration-neural couplings in cortico-basal ganglia circuits, underscoring its potential role in coordinating body-brain interactions during sleep and anesthesia.

**Significance Statement:** Neural oscillations are known to couple with respiration rhythms across various brain areas; however, this coupling in the SNr, a major output hub of the basal ganglia, remained unexplored. Current study fills this gap with several novel findings, revealing state-dependent coupling to respiration rhythms in the SNr and M1 of mice. In particular, the strength of respiration-neural coupling varied across multiple states, including NREM sleep, REM sleep, quiet wakefulness, and anesthesia, and was directly related to the delta power, a hallmark of NREM sleep. These findings provide new insight into how respiratory rhythms interact with cortico-basal ganglia circuits and demonstrate that coupling changes dynamically within and across multiple states.

## Introduction

Breathing is a fundamental physiological rhythm, and growing evidence indicates that neural oscillations across widespread brain regions are coupled to this rhythm. Respiration-neural coupling is well-established in olfactory areas such as the olfactory bulb and piriform cortex, where the coupling arises from sensory inputs via nasal airflow stimulating mechanoreceptors in the olfactory epithelium (Fontanini et al., 2003; Fontanini and Bower, 2005; Grosmaitre et al., 2007; Moberly et al., 2018).

Beyond olfactory structures, respiration-entrained local field potentials (LFPs) are also prominent in the hippocampus across various behavioral states including running, quiet wakefulness, sleep, and anesthesia (Chi et al., 2016; Karalis and Sirota, 2022; Lockmann et al., 2016; Yanovsky et al., 2014; Zhong et al., 2017). Importantly, recent human studies have demonstrated that respiration modulates the temporal structure of sleep-related slow oscillations, sleep spindles, and ripples, suggesting a key role for breathing in coordinating hippocampal-cortical activity during NREM sleep and reinforcing memory consolidation during sleep (Ghibaudo et al., 2024; Schreiner et al., 2023; Schwimmbeck et al., 2025; Sheriff et al., 2024).

These respiration-locked neural activities are not confined to olfactory or hippocampal regions. It has been increasingly recognized as a global phenomenon, observed across diverse brain areas in both rodents and humans (Herrero et al., 2018; Juventin et al., 2023; Karalis et al., 2016; Tort et al., 2018; Zhong et al., 2017). Respiration-coupled rhythms are observed across multiple frequency bands including delta, theta, gamma and sharp wave ripples, and are known to be functionally associated with cognition and memory consolidation (Kluger et al., 2021; Schreiner et al., 2023; Zelano et al., 2016). Despite these widespread effects, many questions remain about how far this coupling extends. In particular, it is still unknown whether subcortical structures like the basal ganglia also exhibit respiration-locked neural dynamics.

The basal ganglia, traditionally associated with motor control, are increasingly recognized for their roles in sleep regulation and maintain anatomical connections with brainstem regions involved in arousal and autonomic control (Foster et al., 2021; Franks, 2008; Liu et al., 2020; McElvain et al., 2021). Its major output nucleus, the substantia nigra pars reticulata (SNr), projects to midbrain and pontomedullary structures that are tightly linked to sleep–wake regulation and respiratory control (Liu et al., 2020; McElvain et al., 2021). Recent findings further demonstrate that the SNr can modulate respiration in a cell-type-specific manner via inhibitory projections to noradrenergic neurons in the locus coeruleus (LC) (Gu et al., 2025). In addition, SNr activity is modulated by catecholamines such as dopamine and noradrenaline, which are key modulators of behavioral state transitions (Wang et al., 2010). Basal ganglia disorders like Parkinson’s disease are often associated with both sleep disturbances and respiratory dysfunction, further indicating the basal ganglia’s involvement in these impairments (Aquino et al., 2022; Baille et al., 2016; De Castro Medeiros et al., 2019; Menza et al., 2010; Miranda et al., 2024; Oliveira et al., 2019; Walker et al., 2024).

Given the growing evidence for the significant role of SNr in respiration and sleep, we investigated respiration-locked activity in the SNr and primary motor cortex (M1). While previous work in rats has reported reduced respiration-neural coupling in the cortex and hippocampus during sleep compared to awake state (Girin et al., 2021), detailed state-dependent analysis in mice, particularly in subcortical regions like the SNr, remains limited. Here, we address this gap by characterizing respiration-locked activity in the SNr and M1 across natural NREM and REM sleep, quiet wakefulness (QW), and anesthesia in mice. Respiration and heart rate were recorded using diaphragm electromyography (EMG) and electrocardiography (ECG), simultaneously with LFPs from the SNr and M1. We focused on respiration coupled rhythms across different states, with particular emphasis on delta-band activity, a hallmark of NREM sleep and a key indicator of sleep quality and deprivation (Brown et al., 2012; Hubbard et al., 2020; Long et al., 2021; Uygun and Basheer, 2022).

## Materials and Methods

### Subjects

This experimental protocol was approved by the Institutional Animal Care and Use Committee of Seton Hall University (Approval No. GB2401). All experimental procedures followed the guidelines outlined in the Guide for the Care and Use of Laboratory Animals by the U.S. National Institutes of Health.

We used a total of 17 adult mice: ten C57BL/6J (Jackson Laboratory, stock #000664; 2 females, 8 males) and seven Gad2-Cre mice (Jackson Laboratory, stock #010802; 3 females, 4 males). All animals were 3-12 months old and weighed 28-32 g. We included Gad2-Cre mice to assess whether their respiratory-related neural patterns differ from those of C57BL/6J mice for future studies. We confirmed that the main findings of this study, state-dependent changes in respiration–neural coupling, were consistent across both genotypes (Figure S1A, B). These findings were also consistent across sexes (Figure S1C, D). The animals were housed in groups of four under a 12-hour light/dark cycle (lights on/off at 7:00 AM/PM) with ad libitum access to food and water. After the implantation of chronic electrodes, mice were housed individually throughout the experimental period.

### Surgical procedures

EMG and ECG electrodes were prepared using 7-stranded, flexible, PFA-coated stainless-steel wire with a 0.001” bare diameter (A-M Systems, Cat. No. 793200). For EMG electrodes, a knot was tied at one end, with a small drop of epoxy applied to create a bulk anchor. About 1 cm from the knot, ∼1 mm of insulation was stripped. For ECG electrodes, insulation of one end of the wire was stripped, then rolled and soldered into a ∼1.5 mm circular metal loop that can work as an anchor.

Mice were anesthetized using a ketamine/xylazine mixture (100 mg/kg for Ketamine and 10mg/kg for Xylazine) and placed in a supine position. Anesthesia was confirmed by the absence of withdrawal reflex upon hind paw pinch, and additional doses were administered as necessary throughout the surgery. Body temperature was maintained at 37-38 °C using a heating pad. A small horizontal skin incision (∼1 cm) was made over the sternum without fully exposing the diaphragm. EMG implantation was done similarly as described in previous research (Hérent et al., 2020). The epoxy-anchored end of the EMG wire was secured to the sternum, and the de-insulated site was positioned beneath the diaphragm by laterally guiding a needle connected to the EMG wire through the rib cage (Figure 1A). The wire was fixed in place by tying a knot on the side that came through the rib. For ECG implantation, a needle connected to the ECG wire was inserted through the sternum and advanced transversely until it exited on the left side of the sternum. The circular metal loop was positioned between the skin and muscle above sternum. Both the EMG and ECG wire ends were tunneled subcutaneously toward the head and later connected to an adaptor (Mill-Max Manufacturing Corp.). The incision was sutured using 5-0 silk sutures.

**Figure 1.**
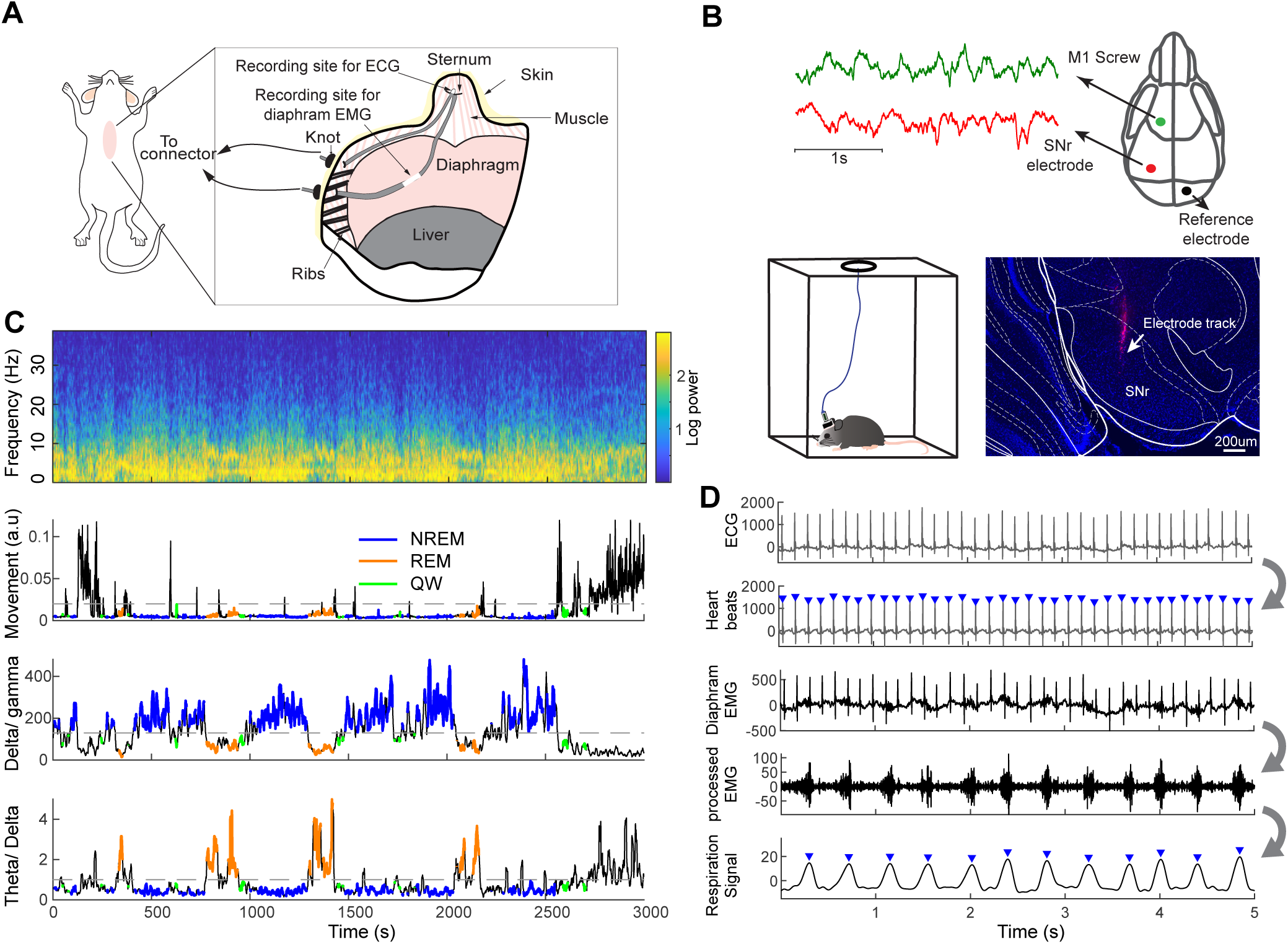
Simultaneous recordings of respiration, heart, and neural signals during sleep. (A) Schematic illustration of chronic electrode Implantation for diaphragm EMG and ECG recordings. (B) Simultaneously with diaphragm EMG and ECG, LFPs were recorded from the M1 using a skull screw and from the SNr using a tungsten electrode. Data was collected in a sleep chamber from freely moving mice. A histological image shows DiI staining of the SNr electrode track. (C) Top row: Spectrogram of M1 LFPs showing changes in delta and theta power across sleep-wake state transitions. Second row: Accelerometer signals were used to monitor movement and identify immobile periods. These periods were classified into NREM sleep, REM sleep and QW states based on delta-to-gamma (third row) and theta-to-delta ratios (bottom row) of the M1 LFPs. (D) Heartbeats were extracted from the ECG signal. The diaphragm EMG was band-pass filtered after removal of heartbeat artifacts (processed EMG). A sliding window calculation of standard deviation was then applied to derive the respiration signal. Inverted blue triangles indicate detected heartbeats and breaths. EMG, electromyogram; ECG, electrocardiogram; LFPs, local field potentials; M1, motor cortex; SNr, substantia nigra pars reticulata; NREM, non-rapid eye movement; REM, rapid eye movement; QW, quiet wakefulness.

Following EMG and ECG implantation, mice were secured in a stereotaxic frame, and small craniotomies (0.5-1.0 mm in diameter) were drilled for SNr LFP electrode and skull screws. Twisted pairs of PFA-coated tungsten wires (0.002” bare, A-M Systems, Cat. No. 792500) were dipped in DiI (DiIC_18_(3), Invitrogen) to mark the electrode tracks for histological verification and implanted in SNr (coordinates: AP: -3.15 mm, ML: -1.5 mm, DV: 4.8 mm from Bregma). Stainless steel screws (Bone Anchor Screws, Stoelting Co.) soldered to tungsten wires were fixed to the skull over the left primary motor cortex (M1, AP: 1.5 mm, ML: -1.5 mm from Bregma) and the right cerebellum (reference). All wires were connected to an adaptor mounted on top of the skull using dental cement.

### Respiratory and electrophysiological recordings

Electrophysiology recordings began 6-7 days post-surgery. Sleep data were collected in a mouse chamber (16 × 16 × 16 cm) across three 4-hour sessions per mouse (Figure1B Bottom) with free access to water. Signals from the M1, SNr, diaphragm EMG, and ECG were recorded simultaneously using an Intan RHD2000 recording system with RHD headstages (Intan Technologies, 0.1-500 Hz bandpass, 1 kHz sampling rate). Movement was detected using the headstage-mounted accelerometer, and mouse behavior was also monitored via video camera to confirm sleep.

In a subset of mice (*n*=14 out of 17), electrophysiology signals were recorded under ketamine/xylazine anesthesia using the same parameters as during sleep recordings. Ketamine (100 mg/kg) and xylazine (10 mg/kg) were administered, and mice were placed on a heating pad maintained at 37°C. Recordings continued uninterrupted for 60-90 minutes until the mice regained consciousness.

### Histology

After completion of data collection, mice were deeply anesthetized and transcardially perfused with phosphate-buffered saline (PBS), followed by 4% paraformaldehyde (PFA). Brains were extracted, post-fixed in 4% PFA at 4 °C for approximately 24 hours, and then transferred to a 30% sucrose solution for cryoprotection until they sank. The brains were sectioned into 50-μm coronal slices using a cryostat (Leica CM1860) and then mounted with a DAPI-containing mounting medium (Fluoromount-G with DAPI, Invitrogen). Electrode tracks labeled with DiI were imaged using a fluorescence microscope (Leica DMi8) or a confocal microscope (Leica Stellaris5) to confirm targeting of SNr. Electrodes missed the SNr in 3 of the 17 mice (2 out of 14 mice for anesthesia recording); thus, their data were excluded from SNr analysis.

## Data analysis

### Behavioral state categorization

To identify periods of QW, NREM sleep, and REM sleep, we used ratios of delta-to-gamma and theta-to-delta band power in conjunction with accelerometer data. Movement was quantified by calculating the standard deviation of accelerometer signals with 1-s sliding window in each axis, then averaging across all three axes. To focus and compare behavioral states without movement, time segments with any movement (threshold of 0.2) were excluded. The M1 signal was then analyzed using the ‘spectrogram’ function in MATLAB, with a 2-s window and a 1-s sliding window.

Power in the delta (0.5-4Hz), theta (5-10Hz), and gamma (30-59Hz) frequency bands were extracted. Time points with high delta-to-gamma (threshold of 130) and low theta-to-delta (threshold of 1) were classified as NREM sleep (Figure 1C). Conversely, low delta-to-gamma and high theta-to-delta ratios were categorized as REM sleep and the remaining segments were labeled as QW state. Sessions with noisy signals or missing any of the three states were excluded (3 out of 51 sessions across 17 mice). Additionally, periods where a single state did not persist for at least 10 seconds were excluded from further analysis.

### Anesthesia state

Time segments between 500-3000s after ketamine/xylazine injection was included in the data analysis. The included segments did not have any movement (accelerometer threshold of 0.2) and showed high delta frequency powers. Anesthetized data were compared to the NREM sleep state in each individual.

### Respiration and heart signal analysis

Diaphragm EMG and ECG signals were processed in MATLAB to extract respiration and heart rhythms (Figure 1D). Heartbeats were identified by applying peak detection function (MATLAB ‘findpeaks’) to the ECG signal after bandpass filtering (10-450Hz). The resulting heartbeat timestamps were used both to calculate heart rate and to remove cardiac artifacts from the diaphragm EMG signal. To eliminate heartbeat contamination in the diaphragm EMG signal, 20-ms segments centered on each heartbeat were replaced with neighboring values, and then bandpass filtered (150-350Hz). This processed EMG signal was applied with a 100-ms sliding window standard deviation calculation and bandpass filtered (1-10Hz) to derive the ‘respiration signal’. Inspiration peaks were identified via a peak detection method (‘findpeaks’) on this respiration signal. To validate peak detection accuracy, 10-s segments were manually reviewed at 5-min intervals across the data, and parameters for ‘findpeaks’ were adjusted as needed to ensure successful detection.

### Spectral power of M1, SNr and respiration signals

M1, SNr and respiration signals were first detrended (MATLAB ‘detrend’). Each signal was then bandpass filtered across 1 to 10Hz range using 1Hz windows with 0.5Hz steps. The Hilbert transform was applied to each band-filtered signal to compute the power at each frequency band. To facilitate peak frequency comparisons across signals, spectral power values were normalized within each session to a 0-1 range using the minimum and maximum power values of each signal.

### Phase locking value (PLV) analysis

To assess phase synchrony between the respiration signal and neural oscillations, PLVs were computed (Figure 2C). Instantaneous phases of the respiration signal and the bandpass (1-5Hz) filtered M1 or SNr signals were obtained using Hilbert transform. Phase differences (Δ (t)) between signal pairs were calculated at each time point (t), and the PLVs were computed as the magnitude of the mean phase difference vector within each 10s window. A 10s window length was chosen because a prior study showed that coupling measures can be sensitive to short window, but segments of 10s or longer yield reliable estimates (Dvorak and Fenton, 2014).

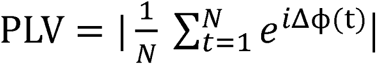

**Figure 2.**
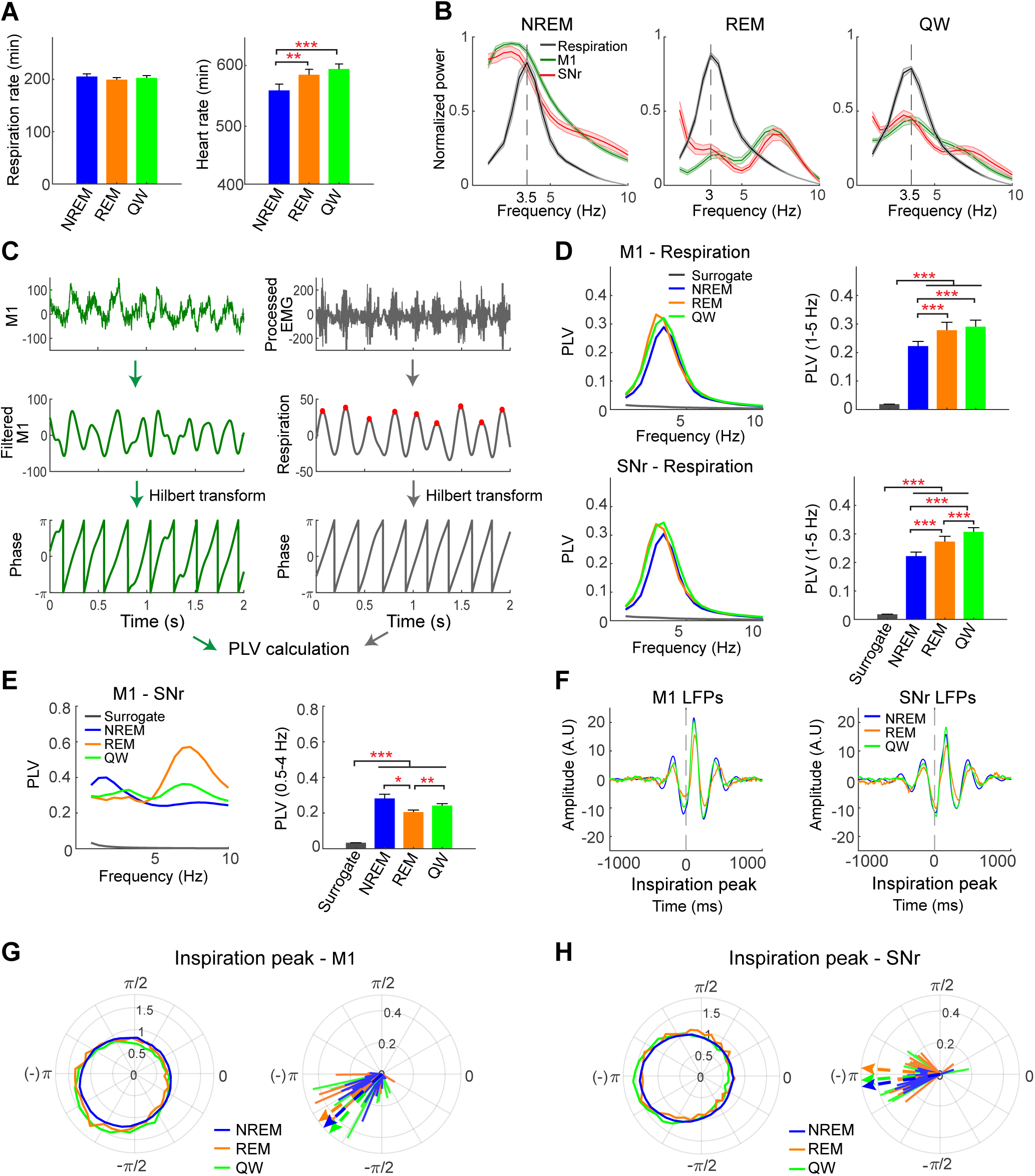
Coupling of M1 and SNr LFPs to respiration rhythms during natural sleep. (A) Left: Respiration rates did not differ significantly across the three states (repeated-measures ANOVA, *F*(2,32) = 1.61, *p*=0.22). Right: Heart rates differed significantly across states (repeated-measures ANOVA, *F*(2,32) =27.6, *p*=1.09×10^-7^), with reduced heart rate during NREM sleep compared to REM and QW (Wilcoxon signed-rank test, *n*=17). (B) Power spectra of M1, SNr and respiration signals were normalized within each mouse to a 0-1 range (minimum to maximum), then averaged across mice. Shaded areas represent the S.E.M. Respiration showed peak power between 3-4 Hz. NREM sleep showed increased delta power and REM sleep showed increased theta power in both M1 and SNr, consistent with state classification criteria. (C) Coupling was assessed by calculating phase-locking values (PLV) between the bandpass-filtered LFPs and respiration signals. Both signals were processed using Hilbert transform to extract phase values for PLV calculation. Red dots indicate inspiration peaks. (D) Top left: PLV spectra between M1 and respiration rhythms are shown for each state. Top right: PLVs between M1 and respiration rhythms in the 1-5Hz range show significant coupling across all three states compared to surrogate data (Wilcoxon signed-rank test, *n*=17 mice). Coupling during NREM sleep was weaker than during REM or QW states (repeated-measures ANOVA, *F*(2,32)=12.7, *p*=8.76 × 10^-5^; post hoc comparisons: Wilcoxon signed-rank test, *n*=17 mice). Bottom: SNr showed similar results to M1, with reduced coupling during NREM compared to other states (repeated-measures ANOVA, *F*(2,26)=65.7, *p*=6.8 × 10^-11^; post hoc comparisons: Wilcoxon signed-rank test, *n*=14 mice). (E) Left: PLVs between M1 and SNr revealed strongest delta-band coupling during NREM sleep and strongest theta-band coupling during REM sleep. Right: Bar graph confirms stronger delta-band coupling during NREM compared to REM sleep (repeated-measures ANOVA, *F*(2,26)=5.38, *p*=0.01; post hoc comparisons: Wilcoxon signed-rank test, *n*=14). Delta band coupling in all states was significantly bigger than in surrogate data (Wilcoxon signed-rank test, *n*=14). (F) M1 and SNr LFPs aligned to inspiration peaks showed delta rhythms, with highest LFPs peaks following the inspiration peak. (G) Left: Phase distributions of inspiration peaks relative to 1-5Hz filtered M1 LFPs were averaged across mice, showing phase preferences near the LFP trough across all states. Right: For each mouse, the resultant vector of the inspiration peak phases is plotted separately for each state. The dotted arrow indicates the mean angle across mice for each state, showing similar phase alignment of inspiration peaks to M1 LFPs across states. (H). Same format as in (G), but for the SNr. In all panels: \**p*<0.05, \*\**p*<0.01, \*\*\**p*<0.001.

Where N is the number of samples within each segment (10,000 samples for 10-s window with 1000Hz sampling rate). PLV ranges from 0 (no phase locking) to 1 (perfect phase locking). The calculated PLVs were averaged across the time points for each behavioral state in each mouse. To assess PLVs across frequencies, M1 and SNr signals were bandpass filtered from 1 to 10Hz range using 1Hz windows with 0.5Hz steps. For statistical comparison, surrogate data were generated by time-reversing the 10s respiration signal, preserving the spectral and amplitude features of the respiration signal while disrupting its temporal relationship with the neural signals.

Additionally, coherence analysis was performed (Figure S2) following approaches used in previous studies (Girin et al., 2021; Tort et al., 2021; Karalis and Sirota, 2022) . Coherence between respiration and M1, as well as between respiration and SNr, was calculated using 10s segments and MATLAB’s ‘mscohere’ function. The resulting coherence values were averaged across time points for each state.

### Phase of inspiration peaks relative to neural signals

To visualize LFPs aligned to inspiration peaks, 2s of LFP segments centered on each inspiration peak were extracted, averaged across mice, and plotted for each behavioral state. For the polar phase plots, the instantaneous phases of bandpass-filtered (1-10Hz) M1 or SNr signals at each inspiration peak were computed using Hilbert transform. Phase distributions were then averaged across mice for each behavioral state (e.g., Figure 2G left) or used to compute a resultant vector for each state in each individual mouse (e.g., Figure 2G right). To compare phase angles between states, pairwise angular differences were computed (‘circ_dist’ from the Circular Statistics Toolbox), and tested with the Wilcoxon signed-rank test.

### Relations between PLV and delta power

Slow (0.5-2Hz)- and fast (2.5-4Hz)-delta power were separately calculated from the M1 and SNr signals using Hilbert transform. Ten-second segments of neural signals were sorted into ten percentile-based bins (from lowest to highest power in 10% increments) based on slow- or fast-delta power. PLVs between the respiration and neural signals were then plotted across bins to illustrate how PLV varied with delta power (Figure 4). To quantify this relationship, correlation coefficients between slow/fast-delta power and PLV were computed for each brain area within each session. A histogram of the resulting correlation coefficient was plotted, and Wilcoxon signed rank tests were performed to assess whether the distribution significantly differed from zero.

### Infraslow fluctuation patterns

To visualize the infraslow fluctuations in the relationship between respiration and neural signals (M1 or SNr), cross-correlograms were computed using MATLAB ‘xcorr’ function on 10s signal segments with a 1s sliding step. To characterize infraslow fluctuations during NREM sleep and anesthesia, time series of respiration rate, heart rates, and PLVs from M1 and SNr were calculated using 10s windows with a 1s sliding step. Only continuous time series longer than 100s were included in the analysis. All signals were converted to percent-change values relative to their session-wise means. Infraslow fluctuations were then visualized by applying Hilbert transform to these percent-change signals after bandpass filtering (0.01-0.1 Hz with 0.005 increasements). Peak power frequencies were identified for each mouse and compared between NREM sleep and anesthesia using Wilcoxon signed rank tests.

### Statistical analyses

Statistical comparisons were performed using MATLAB (MathWorks). Because of limited sample sizes to confirm normality, non-parametric tests (Wilcoxon signed-rank) were used for state-to-state comparisons of respiration rates, heart rates, PLVs, and circular phase analyses, as well as for comparisons against surrogate data or zero. Repeated-measures ANOVA was used for within-subject comparisons across states (NREM, REM, QW, anesthesia), followed by Wilcoxon signed-rank tests for post hoc pairwise analyses. Correlation analyses were applied to assess relationships between delta-band power and PLVs. More detailed information is provided in each figure or in the main text.

## Results

### SNr and M1 oscillations show weaker coupling to respiration during NREM sleep

To examine how brain rhythms are coupled to respiration across different states while minimizing movement-related effects, we first identified immobile periods and classified them into NREM sleep, REM sleep, and QW based on M1 LFPs (Figure 1C). We then analyzed M1 and SNr LFPs for each state in relation to respiration signals that are derived from diaphragm EMG (Figure 1D). We focused on these states because their respiration rates were relatively similar, reducing confounds from large frequency differences across conditions. We excluded the active movement state due to respiration rate differences and the presence of movement-related artifacts in some mice although; however, previous studies have reported robust neural-respiration coupling during movement (Jung et al., 2023; Zhong et al., 2017).

Respiration rates did not significantly differ across states, whereas heart rate was reduced during NREM sleep compared to QW and REM sleep (Figures 2A). Spectral analysis of respiration, M1 LFPs, and SNr LFPs showed partially overlapping patterns between respiration and neural signals. Specifically, the respiration signal exhibited spectral peaks around 3-4 Hz across all three states (Figure 2B), corresponding to the observed respiration frequencies. Both M1 and SNr LFPs showed small spectral peaks near the respiration frequency during REM sleep and QW states, consistent with previous reports (Biskamp et al., 2017; Chi et al., 2016; Tort et al., 2021; Zhong et al., 2017). However, during NREM sleep, this peak was obscured by strong delta power, lacking a distinct spectral component aligned with respiration.

To quantify respiration–neural coupling, we computed phase-locking values (PLVs) between the respiration signal and LFPs from the M1 and SNr (Figure 2C). PLVs were chosen over coherence values to assess phase consistency without the confounding influence of amplitude fluctuations. Across all states, PLVs were significantly higher than those computed from surrogate data (temporally reversed breathing signal), indicating robust phase synchronization between respiration and neural activity regardless of behavioral state (Figure 2D).

However, the strength of this coupling varied systematically across behavioral states. Both M1 and SNr signals showed significantly weaker coupling to respiration during NREM sleep compared to REM sleep and QW (Figure 2D). The finding is consistent with a prior study in rats showing reduced respiration-related coupling in the cortex and hippocampus during slow wave sleep (Girin et al., 2021). To facilitate comparison with previous studies that used coherence analysis, we also computed coherence values and observed similar results as shown in the PLV analysis (Figure S2).

Delta-band synchronization across widespread brain regions, particularly within cortico- thalamic circuits, is a hallmark of NREM sleep, and has been implicated in sensory gating (Brown et al., 2012; Uygun and Basheer, 2022). To further examine inter-regional dynamics, we computed PLVs between M1 and SNr LFPs and found significantly enhanced delta-band M1-SNr coupling during NREM sleep (Figure 2E). This inverse relationship – stronger regional coupling alongside weaker coupling to respiration rhythm during NREM sleep – supports the idea that delta oscillations facilitate inter-regional communication while reducing sensitivity to external rhythmic inputs such as breathing.

We also examined whether heart rhythms were phase-locked to neural rhythms, which could potentially confound the respiration-neural coupling results. The PLVs between heart and neural rhythms were negligible within the 1-5Hz where respiration-neural coupling occurs, indicating no interference in our respiration coupling analysis (Figure S3A-B). Interestingly, however, the heart rhythm was clearly coupled with neural rhythms within the 5-15Hz range corresponding to heart rate frequencies. As visceral rhythms, including gastric, cardiac, and respiratory, are suggested to influence cortical excitability independently (Engelen et al., 2024), this finding highlights dissociable coupling frequencies for respiration and heart rhythms. Although caution is warranted when interpreting these coupling results, since cardiac activity can introduce artifacts into LFP signals via vascular pulsations (Kern et al., 2013; Stam et al., 2023), recent evidence suggests that heartbeat-related pressure pulsations can genuinely modulate neural activity via mechanosensitive ion channels (Jammal Salameh et al., 2024).

### Phase relationships between inspiration and SNr oscillations vary across states

To examine the phase relations between respiration and neural signals, we analyzed the timing of inspiration peaks relative to M1 and SNr oscillations. Inspiration peak aligned LFPs showed that inspiration typically occurred near the trough of the delta rhythm, with the largest LFPs peak following inspiration (Figure 2F). This temporal alignment is consistent with the notion that respiration leads LFP activity modulations, as shown in prior studies using Granger causality (Karalis and Sirota, 2022; Tort et al., 2021).

Next, we examined the phase of M1 and SNr oscillations at which inspiration peaks occurred. Circular phase distribution showed no clear differences between states (Figure 2G, H; Wilcoxon signed-rank test on phase distance between states, NREM vs. REM sleeps in M1, *n*=17, *p*=0.38; SNr, *n*=14, *p*=0.07; NREM vs. QW in M1, *n*=17, *p*=0.19; SNr, *n*=14, *p*=0.03).

### Coupling to respiration under anesthesia increases strongly in SNr but not in M1

Anesthesia is often compared to NREM sleep due to its similarities to deep sleep; however, it also exhibits distinct physiological features (Akeju and Brown, 2017; Franks, 2008), which warrant caution when interpreting studies conducted under anesthetized conditions. Many studies of respiration-neural coupling have been conducted under anesthesia (Brankačk et al., 2025; Fontanini et al., 2003; Juventin et al., 2023; Lockmann et al., 2016; Yanovsky et al., 2014), so direct comparisons with natural sleep states are important for addressing the current gap in understanding the shared and distinct features of respiratory coupling across these states. To enable such comparisons within the same animal, we also recorded respiration, M1, and SNr LFPs under ketamine/xylazine anesthesia in a subset of mice that also underwent natural sleep recordings (*n*=14 out of 17 for M1 and *n*=12 out of 14 for SNr).

During anesthesia, both respiration and heart rates were significantly reduced compared to NREM sleep (Figure 3A), and power spectral analysis of the respiratory signal revealed a clear peak at 2.5 Hz (Figure 3B), reflecting the slowed respiration rate. This was accompanied by a marked increase in delta power (Figure 3B,C), consistent with previous reports (Magill et al., 2004). Both M1 and SNr exhibited strong delta-band activity, with peak power below 2 Hz during anesthesia. Notably, the SNr signals displayed an additional spectral peak near 2.5 Hz, matching the frequency of respiration.

**Figure 3.**
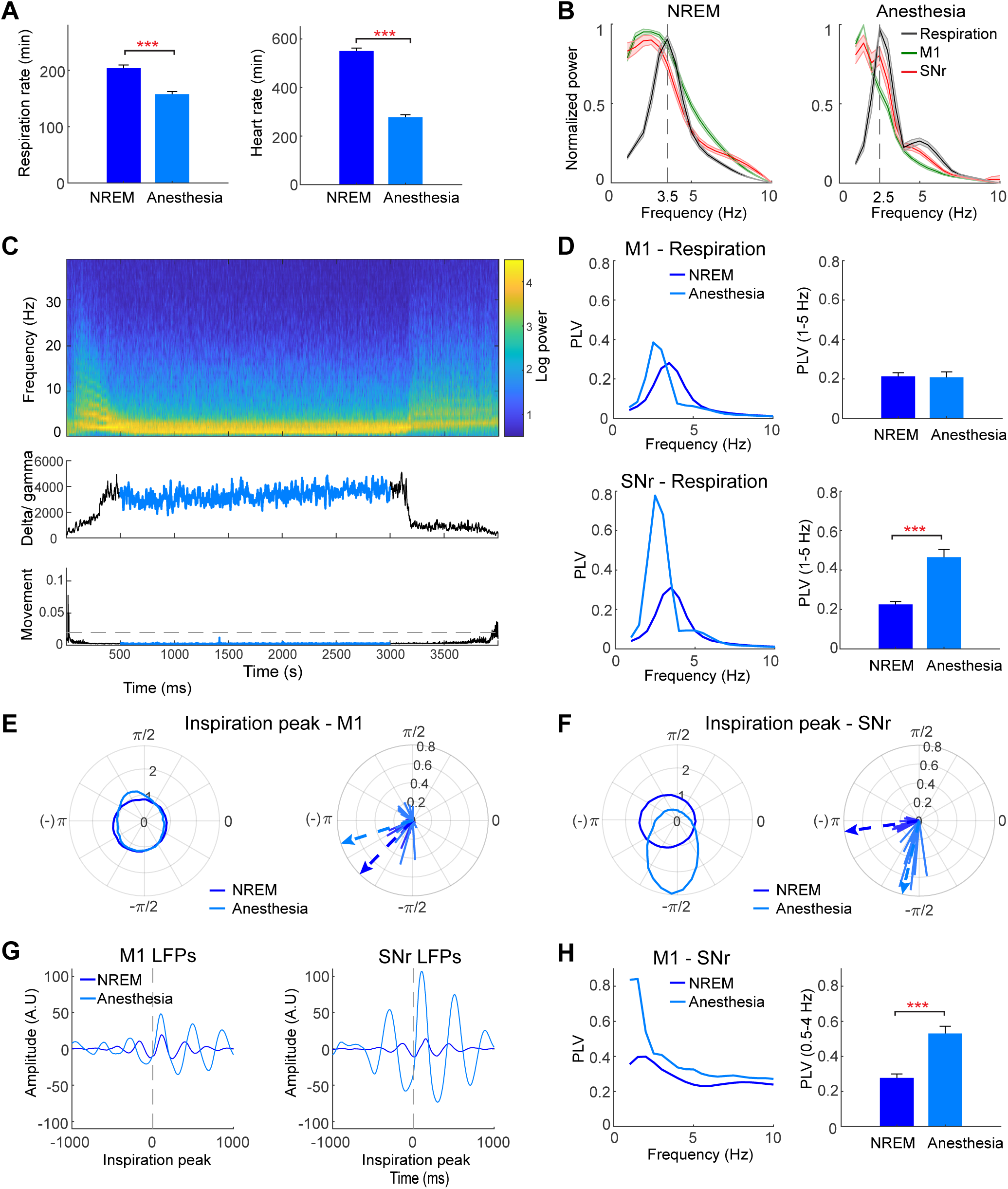
Coupling of M1 and SNr LFPs to respiration rhythms during anesthesia. (A) Both respiration and heart rates were significantly reduced during anesthesia compared to NREM sleep (Wilcoxon signed-rank test, *n*=14 mice). (B) Power spectra of M1, SNr, and respiration signals were normalized within each mouse to a 0-1 range and then averaged across mice. Shaded areas represent the S.E.M. Respiration showed peak power at around 2.5 Hz during anesthesia, and both M1 and SNr exhibited strong delta power. (C) Example M1 LFP spectrogram (top) shows a marked increase in delta power following ketamine/xylazine administration (time 0). Signals from 500 to 3000s post-injection were analyzed, during which all recorded mice showed sustained delta elevation (middle) and no movement (bottom). (D) Spectral peaks of M1-respiraton and SNr-respiration PLVs are shifted to lower frequencies during anesthesia, consistent with slower respiration rates compared to NREM sleep (top and bottom left). Bar graphs of PLVs in the 1-5Hz range show significantly increased coupling to respiration in the SNr, but not in M1, during anesthesia (top and bottom right, Wilcoxon signed-rank test, *n*=14 mice for M1 and *n*=12 mice for SNr). (E-F) Same format as in Figure 2(G-H), but for comparing NREM sleep and anesthesia. The phase of inspiration peaks relative to M1 LFPs did not differ significantly between NREM sleep and anesthesia (E), whereas in SNr, inspiration peaks positioned at significantly shifted phase angles and showed stronger phase locking during anesthesia (F). (G) M1 and SNr LFPs aligned to inspiration peaks. SNr LFPs showed prominent delta rhythms during anesthesia, reflecting both elevated delta power and stronger respiration coupling. (H). Left: PLVs between M1 and SNr showed increased delta-range coupling during anesthesia, particularly in the slow delta band (<2Hz). Right: The bar graph confirms a significant increase in M1-SNr delta coupling during anesthesia compared to NREM sleep. In all panels: \*\*\**p*<0.001.

PLV analysis of respiration-neural coupling showed that the peak frequency of coupling shifted to lower frequencies under anesthesia compared to NREM sleep, consistent with the reduced respiration rate (Figure. 3D, left). Coupling power in the 1-5Hz band was similar in M1 between NREM sleep and anesthesia (Figure 3D, top right); however, SNr-respiration coupling was dramatically enhanced during anesthesia (Figure 3D, bottom). Circular phase distributions and inspiration peak-aligned LFPs also showed stronger and more consistent coupling between respiration and SNr LFPs under anesthesia (Figure 3E-G). In addition, while the phase of inspiration peaks in M1 did not differ significantly between the two states, the phase in SNr showed a significant shift (Wilcoxon signed-rank test on phase distance between NREM and anesthesia; M1: *n*=14, *p*=0.46; SNr: *n*=12, *p*=0.0004). Given the limited knowledge about SNr-respiration coupling and the absence of known direct olfactory inputs to the SNr, the marked increase in SNr-respiration coupling under anesthesia represents a novel and unexpected finding.

Spectral PLV analysis between M1 and SNr signals revealed a stronger peak in the delta range during anesthesia compared to NREM sleep (Figure 3H, left), confirmed by statistical comparison of PLVs in the delta (0.5-4 Hz) band (Figure 3H, right). When the delta band was further subdivided into slow (0.5-2Hz)- and fast (2.5-4Hz)-delta, only the slow-delta band showed a significant increase in the M1-SNr coupling under anesthesia (*n*=12, Wilcoxon signed-rank test, *p*=4.8×10^-4^), while the fast-delta band did not (*p*=0.05). This enhanced synchronization in the slow-delta range suggests its specific role in cortico-basal ganglia communication during anesthesia. Notably, this slow-delta frequency range was separable from the peak frequency of respiration-neural coupling, which was near the fast-delta band. This separation indicates a systematical dissociation between inter-regional communication and respiration-neural couplings.

Heart rates were significantly reduced during anesthesia and partially overlapping with respiration frequencies. Consequently, PLV analysis between heart and neural rhythms during anesthesia showed enhanced coupling within the 1-5 Hz range compared to NREM sleep (Figure S3C, D). This raises the possibility that the observed enhancement in SNr-respiration coupling during anesthesia could be partially contaminated by cardiac activity. However, heart-neural coupling did not show region-specific increase as we observed in the SNr-respiration coupling. Moreover, the magnitude of SNr-heart coupling (<0.1) was negligible compared to SNr-respiration coupling (around 0.5), minimizing the likelihood of cardiac contamination in our respiration-related findings.

The ketamine-xylazine mixture used for anesthesia acts on multiple targets including NMDA receptors and α₂-adrenoceptors, making it difficult to identify the exact mechanism underlying the SNr-specific increase in coupling. To investigate this further, we tested a subset of mice (*n* = 6) using ketamine alone at the same dose (100 mg/kg). The resulting coupling pattern under ketamine monotherapy resembled that of NREM sleep rather than that of ketamine-xylazine anesthesia (Figure S4), indicating that ketamine alone is insufficient to account for the strong SNr-respiration coupling observed with the mixture. We did not test xylazine monotherapy, as it can induce severe and unpredictable adverse effects, including bradycardia, hypotension, and respiratory depression.

### Slow and fast delta rhythms show opposing relationships to respiration-neural coupling

Slow-delta rhythm is a hallmark of deep NREM sleep and known to be associated with stronger suppression of sensory inputs. Distinct features between slow- and fast-delta rhythms have been also reported under conditions such as sleep deprivation (Hubbard et al., 2020). To further investigate how these delta sub-bands relate to respiration- neural coupling, we analyzed the relationship between delta power and coupling strength during NREM sleep (Figure 4A-C). We divided slow-delta (0.5-2 Hz) and fast-delta (2.5-4Hz) power into 10 percentile bins. PLVs were then calculated separately for data segments within each bin, and the resulting values were visualized (Figure 4B,C left). Overall, respiration-neural coupling decreased with stronger slow-delta power, whereas it increased with stronger fast-delta power. This opposite impact of slow- and fast-delta rhythms on coupling was observed in both M1 and SNr. The results were confirmed by significant negative and positive correlations between PLV values and slow- and fast-delta power, respectively (Figure 4B,C right).

Under ketamine/xylazine anesthesia (Figure 4D), similar trends were observed in the M1: slow-delta power was negatively associated with respiration-M1 coupling, while fast-delta power showed a positive relationship (Figure 4E,F top). However, the SNr did not exhibit a significant relationship between slow-delta power and respiration coupling, although the positive association with fast-delta power remained particularly strong (Fig. 4E, F bottom).

These findings suggest that slow-delta oscillations may suppress respiration-related synchronization, while fast-delta rhythms enhance it. Furthermore, the SNr appears less sensitive to slow-delta modulation under anesthesia, highlighting differences between cortical and subcortical contributions to respiratory coupling dynamics.

### Respiration-neural coupling shows fluctuating patterns in Infraslow timescale

We showed the state-dependent changes in respiration-neural coupling; however, it remains unclear whether this coupling is maintained stable within each state or exhibits dynamic fluctuations. To investigate this, we examined respiration-neural coupling over a long timescale during natural sleep and anesthesia, alongside other physiological parameters such as respiration and heart rates. Representative examples of natural sleep showed that the coupling (PLVs) between respiration and M1/SNr signals fluctuated with infraslow cycles on the order of tens of seconds, and cross-correlograms confirmed time-varying phase alignment with respiration (Figure 5A). Other physiological parameters (respiratory and heart rate) also exhibited infraslow dynamics. Under anesthesia (Figure 5B), infraslow fluctuations of respiration-neural coupling persisted; however, both respiration and heart rates show more stable, less variable patterns.

**Figure 4.**
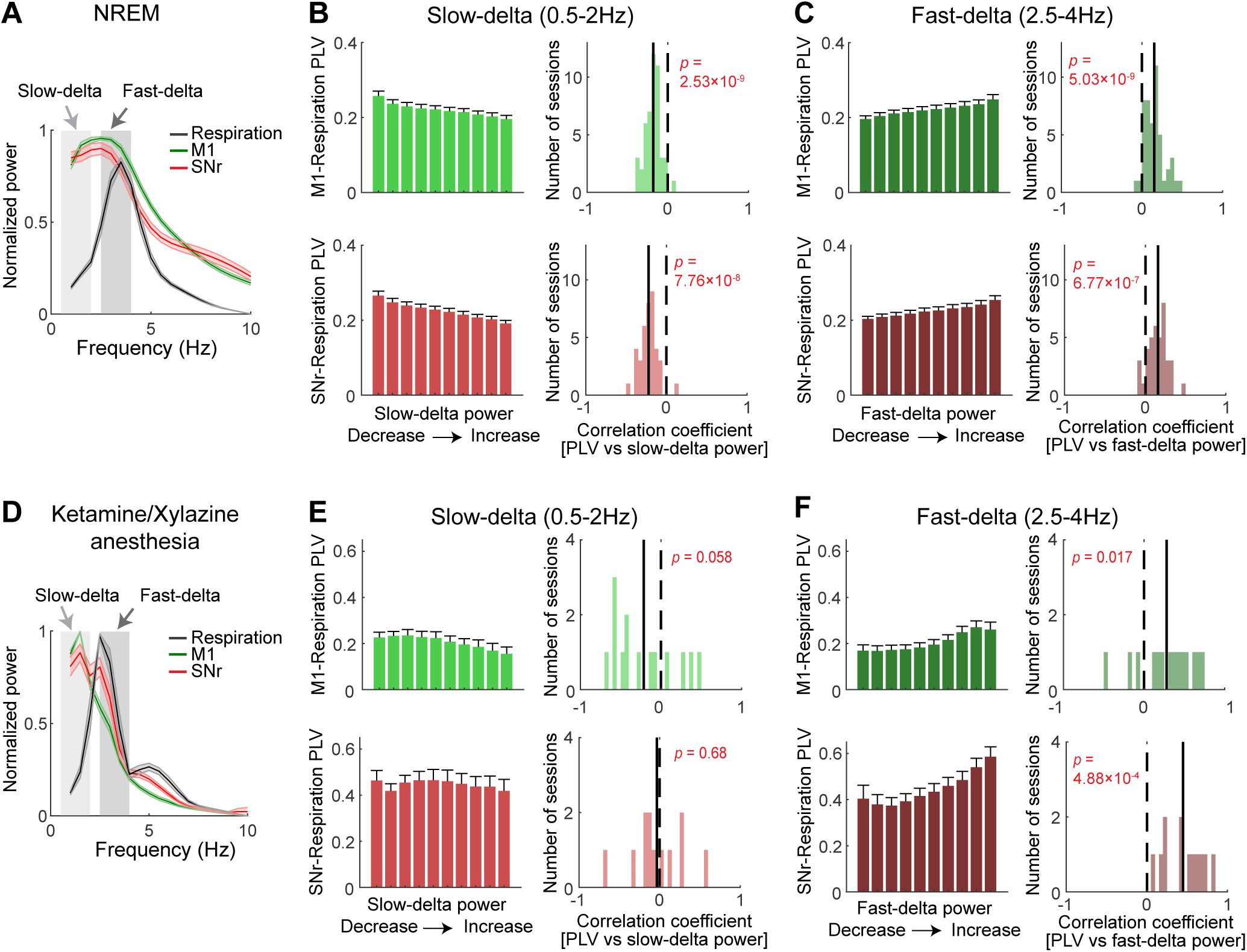
Relationship between delta power and respiratory coupling. (A) Time series of slow (0.5-2Hz) and fast (2.5-4Hz) delta power from M1 and SNr LFPs were extracted during NREM sleep. (B) Top left: Time points were categorized into low and high slow- delta power in M1, and corresponding M1-respiration PLVs were plotted. Higher slow- delta power was associated with reduced M1-respiration coupling. Top right: Correlation coefficients between PLVs and slow-delta power in M1 were significantly shifted toward negative values (Wilcoxon signed-rank test, *n*=48 sessions). Black line indicates mean value across sessions and dotted line marks zero. Bottom left: A similar pattern was observed between slow-delta power and SNr-respiration PLVs. Bottom right: In SNr, PLVs were also negatively correlated with slow-delta power (Wilcoxon signed-rank test, *n*=41 sessions). (C) Coupling to respiration increased with higher fast-delta power in both M1 and SNr. Same format as in (B). (D) Slow- and fast-delta powers from M1 and SNr were extracted during anesthesia. (E-F) During anesthesia, M1-respiration coupling showed similar relationships with delta power (Wilcoxon signed-rank test, *n*=14 sessions) as observed during NREM sleep. In contrast, SNr-respiration coupling showed no significant association with slow-delta power, but a strong positive correlation with fast-delta power (Wilcoxon signed-rank test, *n*=12 sessions). Same format as in (B-C).

**Figure 5.**
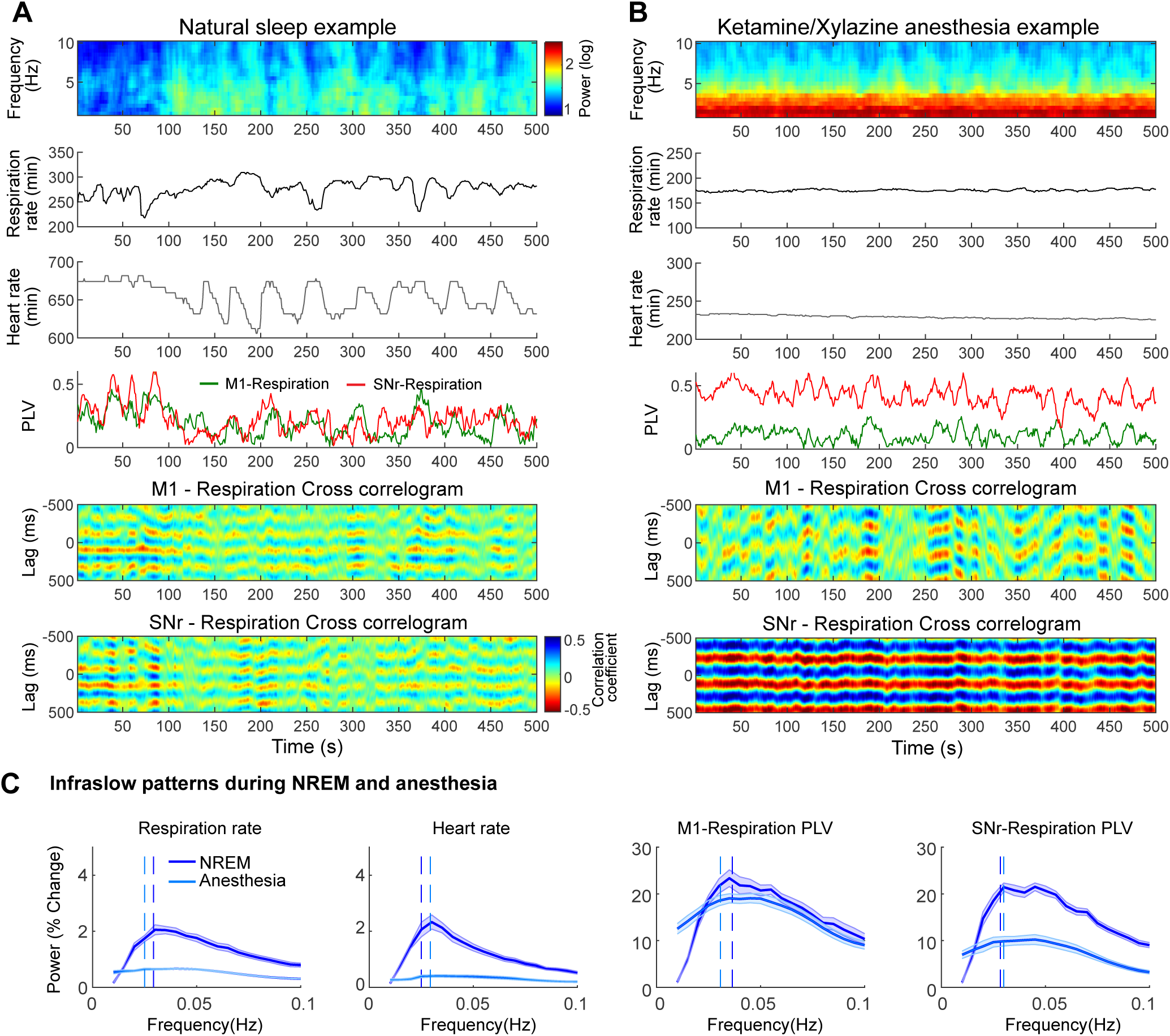
Infraslow fluctuations of physiological and neural signals during sleep and anesthesia. (A-B) Representative examples from a single mouse during natural sleep- wake cycles (A) and ketamine/xylazine anesthesia (B). From top to bottom: Spectrogram of M1 LFP power (1-10Hz), respiration rates, heart rates, PLVs between M1/SNr LFPs and respiration signals, and cross-correlograms (respiration-M1 LFPs and respiration-SNr LFPs). To visualize the temporal dynamics of respiration-neural coupling, cross-correlograms were calculated using 10s window segment with a 1s sliding window. Strong color contrast (high correlation coefficient) indicates stronger respiratory coupling. All signals show slow fluctuations during natural sleep, while respiration and heart rates remain stable under anesthesia. (C) To illustrate infraslow fluctuations (0.01–0.1 Hz) during NREM sleep and anesthesia, group-averaged power spectra of the measured signals (normalized as % change) are plotted (*n*=14 mice for all signals except SNr- respiration PLV, *n*=12 mice). Shaded areas represent ± SEM. Vertical dashed lines indicate mean peak frequency, which was consistently observed around 0.03 Hz across most signals.

To examine the spectral properties of these infraslow (0.01-0.1 Hz) fluctuations, each signal (respiration rate, heart rate, and respiration-neural coupling) was normalized as percent-change, and their spectral power distributions were visualized. The mean spectral peak frequency was approximately 0.03Hz in all signals (Figure 5C). Comparison of peak frequencies between NREM sleep and anesthesia revealed no significant differences (*n*=14, Wilcoxon signed-rank test, *p*>0.05 in all signals), suggesting the presence of a shared infraslow modulating mechanism across both states. However, as shown in the example, respiration and heart rates during anesthesia were more stable and persistent over time, resulting in lower percent-change power and near-flat spectral patterns compared to NREM sleep.

## Discussion

In this study, we provide the first detailed characterization of respiration–neural coupling across multiple states—including QW, NREM sleep, REM sleep, and anesthesia—in the SNr and M1, two regions not previously studied in this context. Across all states, both structures exhibited respiration-locked activity, but coupling strength and phase characteristics varied by state. Notably, coupling dynamics were tightly linked to slow- and fast-delta power, revealing a previously unrecognized interaction between cortico-basal ganglia circuit and respiratory rhythms.

Previous research on respiratory coupling has largely focused on olfaction-related regions, hippocampus, frontal and parietal cortex, and sensory areas; however, studies investigating motor cortex and basal ganglia have been limited. While a previous study reported respiration-related rhythms in the motor cortex of rats during exploration, it did not assess state-dependent changes (Rojas-Líbano et al., 2018). In addition, to our knowledge, no previous study has demonstrated respiratory coupling in the basal ganglia. Our study addresses this gap by demonstrating state-dependent respiration- neural couplings in the motor cortex and SNr. Notably, emerging evidence supports that specific SNr cell types can regulate both sleep-wake cycles and respiration rates (Liu et al., 2020; Gu et al., 2025), highlighting the SNr’s critical role in these processes. Our current study adds to this evolving picture by providing evidence that the SNr is not only involved in regulating respiration rates but is also synchronized with respiration rhythms.

Coupling strength was significantly attenuated during NREM sleep compared to QW and REM sleep in both M1 and SNr. This finding aligns with other studies in rats and cats, which showed diminished respiration-coupled neural signals during NREM sleep relative to wakefulness (Cavelli et al., 2020; Girin et al., 2021). However, despite this attenuation, respiration-neural coupling remained robustly above surrogate levels during NREM sleep, suggesting its continued functional significance. Supporting this, recent human studies have demonstrated that respiration rhythms during NREM sleep coordinate sleep-related oscillations such as sleep spindles, ripple, and slow oscillations, thereby contributing to memory consolidation (Schreiner et al., 2023; Ghibaudo et al., 2024; Sheriff et al., 2024; Schwimmbeck et al., 2025).

To better understand the significance of respiration-neural coupling during sleep, we further examined its relationships with delta band power, a hallmark of NREM sleep, by sub-dividing the band into slow-delta (0.5-2 Hz) and fast-delta (2.5-4 Hz) components.

We found that slow-delta power was negatively correlated with the strength of respiration-neural coupling, suggesting that increases in slow-delta activity contribute to the attenuation of respiration-coupling during NREM sleep. Slow-delta rhythms are known to be associated with deeper stages of NREM sleep and are implicated in long-range inter-regional communication (Magill et al., 2004; Hubbard et al., 2020; Uygun and Basheer, 2022). Consistent with this, we found that regional coupling between M1 and SNr was most prominent in this slow-delta frequency range. Together, these findings suggest that global neural synchronization in slow-delta rhythms during deep sleep attenuates the responsiveness of cortico-basal ganglia circuits to respiration rhythms.

On the other hand, fast-delta power during NREM sleep was positively correlated with the strength of respiration-neural coupling. Fast-delta, also called as δ2, has been reported to be more sensitive to sleep deprivation, and its dynamic features resemble physiological changes such as temperature, muscle tone and heart rate (Hubbard et al., 2020). However, the fast-delta frequency range overlaps with respiration frequency in mice, making it difficult to determine whether pre-existing sleep-related delta rhythms are coupled to respiration, or they are simply co-existing with respiration-related rhythms. In other species with slower respiration rates such as rats (1-2Hz) and humans (0.2-0.3Hz), respiration-related neural rhythms occur at the corresponding respiration frequency during NREM sleep (Girin et al., 2021). However, delta waves during NREM sleep emerge within a similar frequency range (0.5-4 Hz) across species, and the overlap between fast-delta and respiration frequency appears to be unique in mice. Thus, our findings from mice should be interpreted with caution when generalizing to other species. Comparative studies are needed to clarify how delta sub-bands interact with respiratory rhythms and to uncover species-specific mechanisms. Causal approaches such as optogenetic or chemogenetic manipulations could further test delta-respiration relationships, though selectively targeting distributed delta-generating circuits remains technically challenging.

During ketamine/xylazine anesthesia, SNr LFPs revealed two spectral peaks—one reflecting slow-delta, and another in fast-delta, the latter overlapping with the respiration frequency. This dual-peak pattern in the SNr resembles findings in the hippocampus of urethane-anesthetized rats, which showed the presence of both intrinsic slow oscillation (<1.5Hz) and respiration-related rhythm (Lockmann et al., 2016). These two rhythms are thought to have different origins as evidenced by their distinctive features – slow oscillations are coupled to neocortical up-and-down transitions, and respiration-related rhythm are coupled to olfactory bulb rhythms and disappeared with tracheotomy. Similarly, our results show strong inter-regional coupling between M1 and SNr in slow-delta bands and respiration-locked activity in fast-delta bands. This dissociation highlights how anesthesia unmasks parallel delta-band processes—one coordinating cortico–basal ganglia dynamics, the other entraining neural activity to peripheral rhythms like respiration.

In the fast-delta range, coupling of SNr to the respiration rhythm dramatically increased during anesthesia compared to NREM sleep, while M1 showed similar coupling strength between the two states. Moreover, circular distribution analysis revealed clearly shifted SNr phase to the inspiration peak during anesthesia compared to NREM sleep, while M1 phase did not change between states. This dissociation indicates a distinct involvement of SNr in NREM sleep and anesthesia. While both states share many features such as thalamocortical deactivation, they are not identical (Brankačk et al., 2025; Franks, 2008), and our findings show the clear differences in the SNr between these two states.

The mechanisms underlying respiration–neural coupling in distant brain regions remain under debate. While nasal airflow and olfactory input is known as the primary contribution to respiration-entrained rhythms, recent studies reported evidence for another source of the coupling, involving non-olfactory. Entrainment of neural firings to respiration rhythms was reported in multiple brain areas even after olfactory deafferentation, and corollary discharges from brainstem respiratory generators were suggested as the non-olfactory mechanisms for this entrainment (Karalis and Sirota, 2022). Basal ganglia output, SNr has diverse connections with the brainstem, including the preBötzinger complex and LC (Yang et al., 2020; McElvain et al., 2021; Gu et al., 2025), key nodes for respiratory rhythm generation and regulating sleep and autonomic functions, respectively. These anatomical connections between SNr and brainstem suggest that their strong coupling to respiration during anesthesia might be mediated by the corollary discharges from brainstem. This mechanism could explain the dissociation between SNr and M1 observed under anesthesia.

Interestingly, respiration-neural coupling was not stable within each state, but instead fluctuated dynamically on an infraslow timescale during both NREM sleep and anesthesia. Recent studies have highlighted the significance of infraslow fluctuations (∼0.02Hz) in physiological and neural signals during natural sleep (Lecci et al., 2017; Bueno-Junior et al., 2023). In particular, sigma power (10-15Hz) showed strong infraslow fluctuations and temporally associated with heart rate fluctuations during NREM sleep in both mice and humans, pointing to a functional integration of cardiovascular and neural rhythms. Our findings showing significant heart-neural coupling within 5-15Hz aligns with these studies. These fluctuations are thought to reflect a dynamic balance between maintaining sensory reactivity to the environment and promoting brain recovery with memory consolidation. Moreover, LC activity has been reported as a key generator of infraslow rhythms, coordinating arousal, autonomic output, and neural excitability (Osorio-Forero et al., 2021, 2025). This framework offers a valuable perspective for interpreting our findings on the temporal dynamics of respiration-neural coupling: as sensory reactivity to the external environment varies with infraslow rhythms during sleep, neural responsiveness to respiratory rhythm fluctuates on a similar time scale. This raises an important question for future study regarding the direct relationship between respiration-neural coupling dynamics and LC activities.

Recently, other infraslow rhythms, such as the gastric basal rhythm (∼0.05Hz), have been identified as potential modulators of brain activity (Rebollo et al., 2018; Rebollo and Tallon-Baudry, 2022; Richter et al., 2017). Gastric rhythms were coupled to the amplitude of cortical alpha rhythms (10-11Hz), and coupling between gastric rhythms and brain activity measured using fMRI was observed across multiple brain areas, especially in sensory and motor cortices. Although gastric activity was not measured in the present study, it is conceivable that similar visceral rhythms in mice could contribute to the infraslow modulations we observed in respiration-neural coupling. Future studies on infraslow rhythms across multiple organs, including the brain, heart, and stomach, will open a new perspective on brain-body interactions.

Collectively, our study reveals state-dependent respiration–neural coupling in the cortico-basal ganglia circuits and demonstrate how these dynamics interact with delta sub-band activity. These findings provide new insights into how internal brain states interact with peripheral rhythms like respiration, with important functional implications for both sleep and anesthesia. Furthermore, elucidating the mechanisms underlying respiration–neural coupling, especially within basal ganglia circuits will shed light on the pathophysiology of conditions such as Parkinson’s disease, where both sleep and respiration are commonly disrupted (Menza et al., 2010; Baille et al., 2016; Aquino et al., 2022; Walker et al., 2024).

