## Supplemental Figure Legend for "Dynamic respiration-neural coupling in substantia nigra across sleep and anesthesia"

**Figure S1.** Respiration-M1/SNr coupling across genotypes and sexes in multiple behavioral states. All groups consistently showed reduced respiration-neural coupling during NREM compared to REM and QW in both M1 and SNr (top), and a marked increase in coupling in the SNr during anesthesia (bottom). (A) C57BL/6J mice: *n*=10 for M1, *n*=8 for SNr. (B) Gad2-Cre mice: *n*=7 for M1, *n*=6 for SNr. (C) Male mice: n=12 for M1, *n*=9 for SNr. (D) Female mice: *n*=5 for both M1 and SNr. Same format as in Figure 2D and 3D. In all panels: **p*<0.05, ***p*<0.01, ****p*<0.001; Wilcoxon signed-rank test.

**Figure S2.** Coherence between respiration and M1/SNr LFPs during sleep and anesthesia. (A) Coherence analysis, instead of PLV analysis, was used to compare NREM sleep, REM sleep, and QW. Same format as in Figure 2D. (B) Coherence analysis comparing NREM sleep and anesthesia. Same format as in Figure 3D. In all panels: **p*<0.05, ***p*<0.01, ***p<0.001.

**Figure S3.** Coupling of M1 and SNr LFPs to heart rhythms across multiple states using analyses similar to those applied to respiration-neural coupling in Figures 2B, D and Figure 3B, D. (A) Normalized power spectra of heart signals, M1 LFPs, and SNr LFPs during NREM, REM and QW states (*n*=17 for heart and M1; *n*=14 for SNr). Peak heart rhythm power occurred around 10 Hz. Vertical dashed lines indicate the peak frequency of the heart rhythm. Same format as in Figure 2B. (B) Coupling between heart and neural rhythms across NREM, REM and QW states showed little coupling at the respiration frequency (1-5Hz, middle) and notable coupling at the heart frequency (5-15Hz, right). Left and middle: same format as in Figure 2D, but using the heart rhythm. PLVs showed no significant differences across behavioral states within the 1-5Hz range (repeated-measures ANOVA; M1, *F*(2,32)=1.12, *p*=0.33; SNr, *F*(2,26)=1.78, *p*=0.19). Within the 5-15Hz, SNr-heart PLVs showed borderline state differences (M1, *F*(2,32)=3.32, *p*=0.049; post hoc comparisons: Wilcoxon signed-rank tests, *n*=17 mice; SNr, *F*(2,26)=3.30, *p*=0.05). (C) Normalized power spectra of heart signals, M1 LFPs, and SNr LFPs during NREM and anesthesia (*n*=14 for heart and M1; *n*=12 for SNr). Peak heart rhythm power occurred between 4-5 Hz during anesthesia. Same format as in Figure 3B but for heart rhythm. (D) Left and middle: While neural coupling to respiration increased markedly in SNr but not in M1, neural coupling to heart rhythm increased in both SNr and M1 due to the reduced heart frequency under anesthesia. Same format as in Figure 3D. Right: No significant differences were observed in 5-15Hz PLVs between heart and neural rhythms. In all panels: **p*<0.05, ****p*<0.001.

**Figure S4.** Comparisons among NREM sleep, ketamine/xylazine anesthesia, and ketamine-only states in a subset of mice (*n*=6). (A) Respiration (left) and heart rate (right) during ketamine-only condition showed patterns more similar to NREM sleep than ketamine/xylazine (K/X) anesthesia. (B) Normalized power spectra of respiration, M1, and SNr signals during NREM sleep, K/X anesthesia, and ketamine-only conditions. Shaded areas represent the S.E.M. (C) PLV between M1 and respiration in the 1–5 Hz range showed no significant differences across NREM, K/X, or ketamine-only states. (D) SNr-respiration PLV was significantly increased under K/X anesthesia compared to both NREM and ketamine-only conditions. (E) M1-SNr coupling in the 0.5-4 Hz range was enhanced during K/X anesthesia compared to NREM sleep. same format as in Figure 3A, B, D, and H. In all panels: Wilcoxon signed-rank test, *n* = 6 mice; **p*<0.05.
