## Supplementary figures and images for "Dynamic respiration-neural coupling in substantia nigra across sleep and anesthesia"

### Supplemental Figure 1

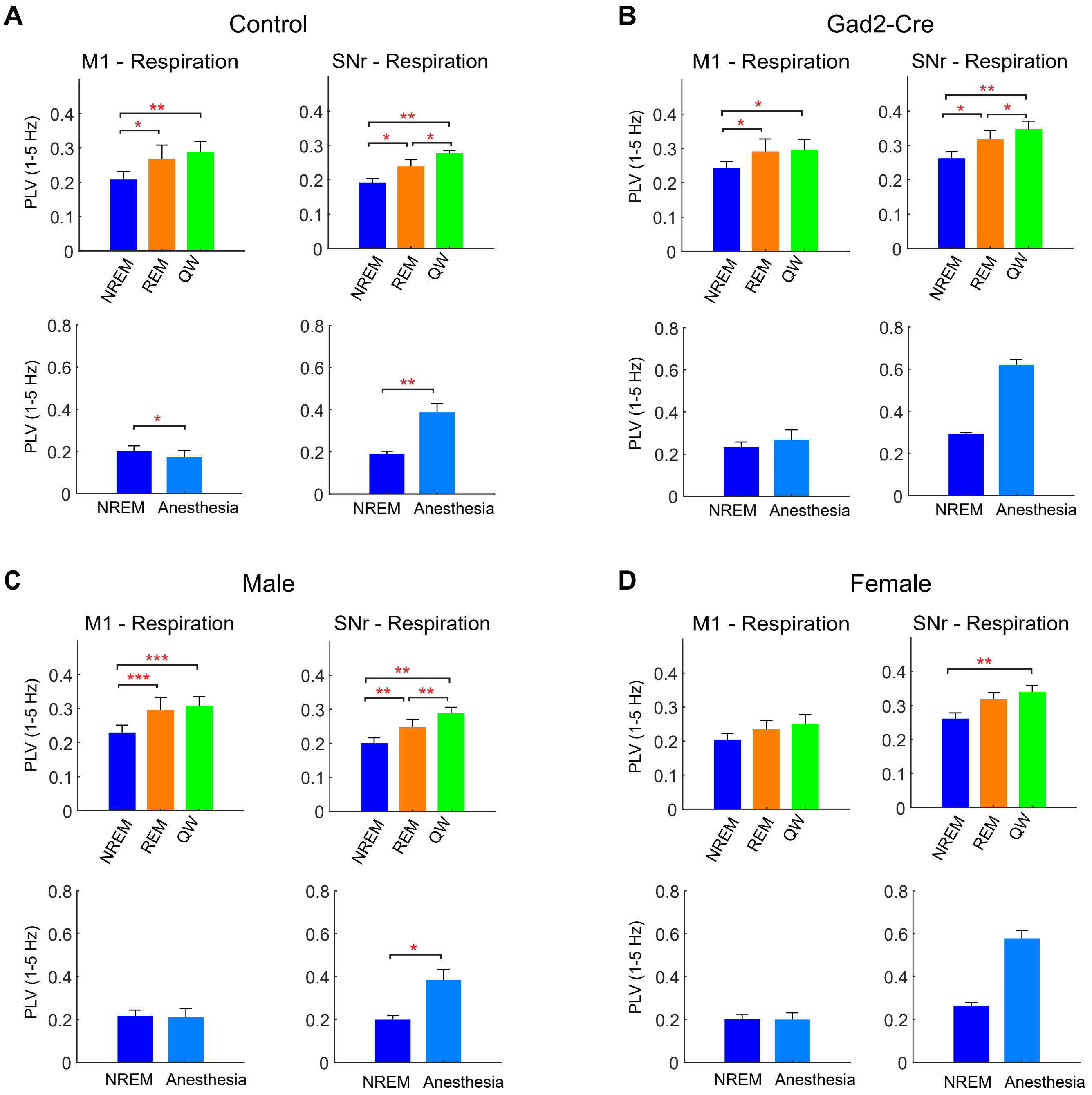

### Supplemental Figure 2

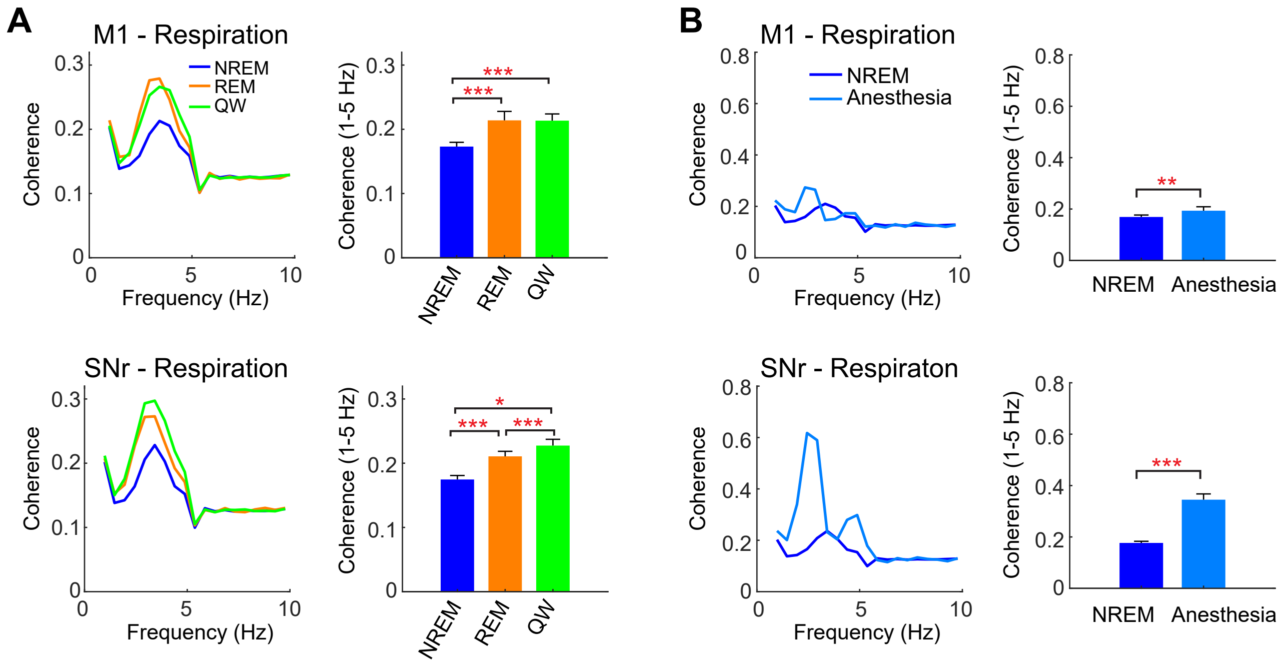

### Supplemental Figure 3

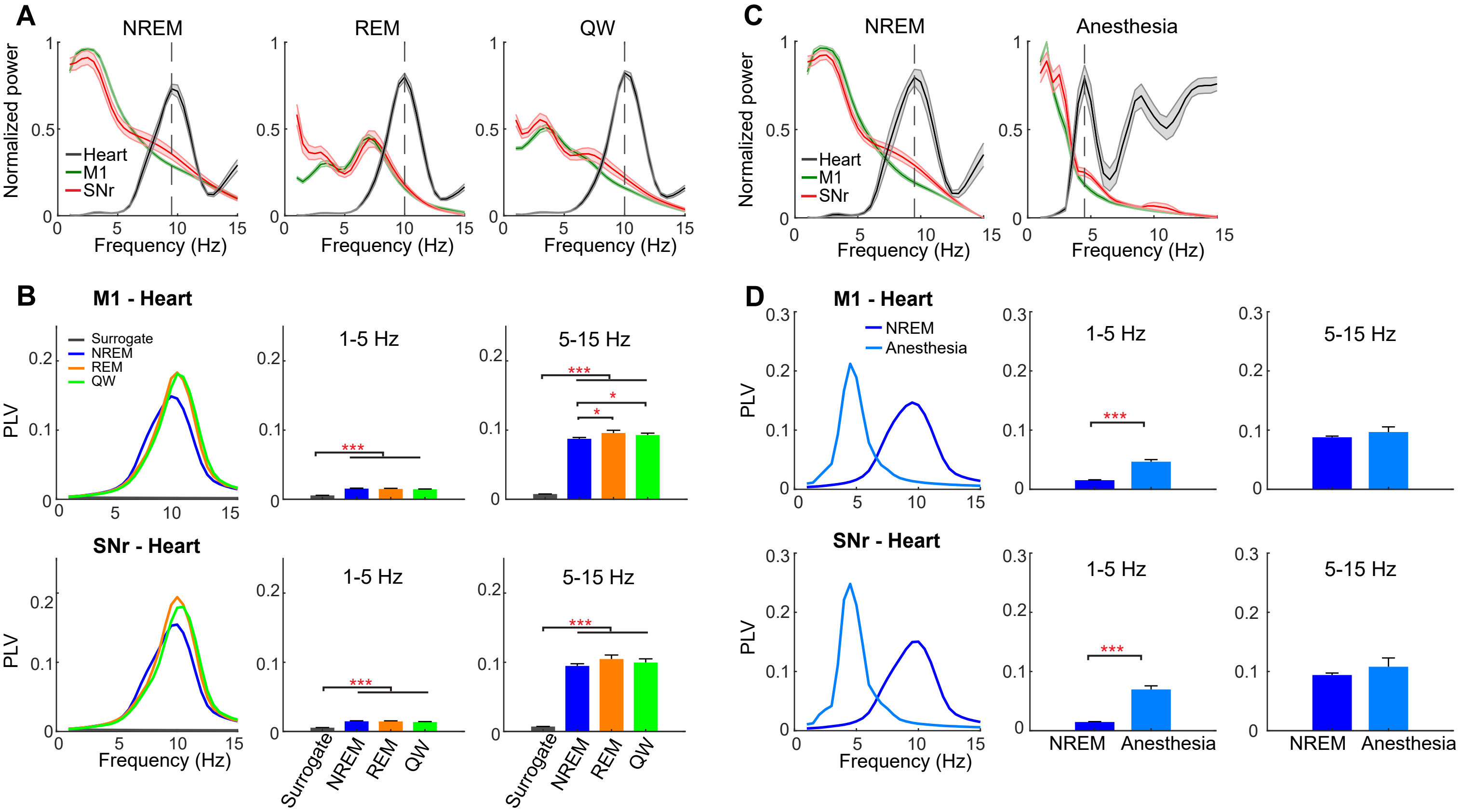

### Supplemental Figure 4

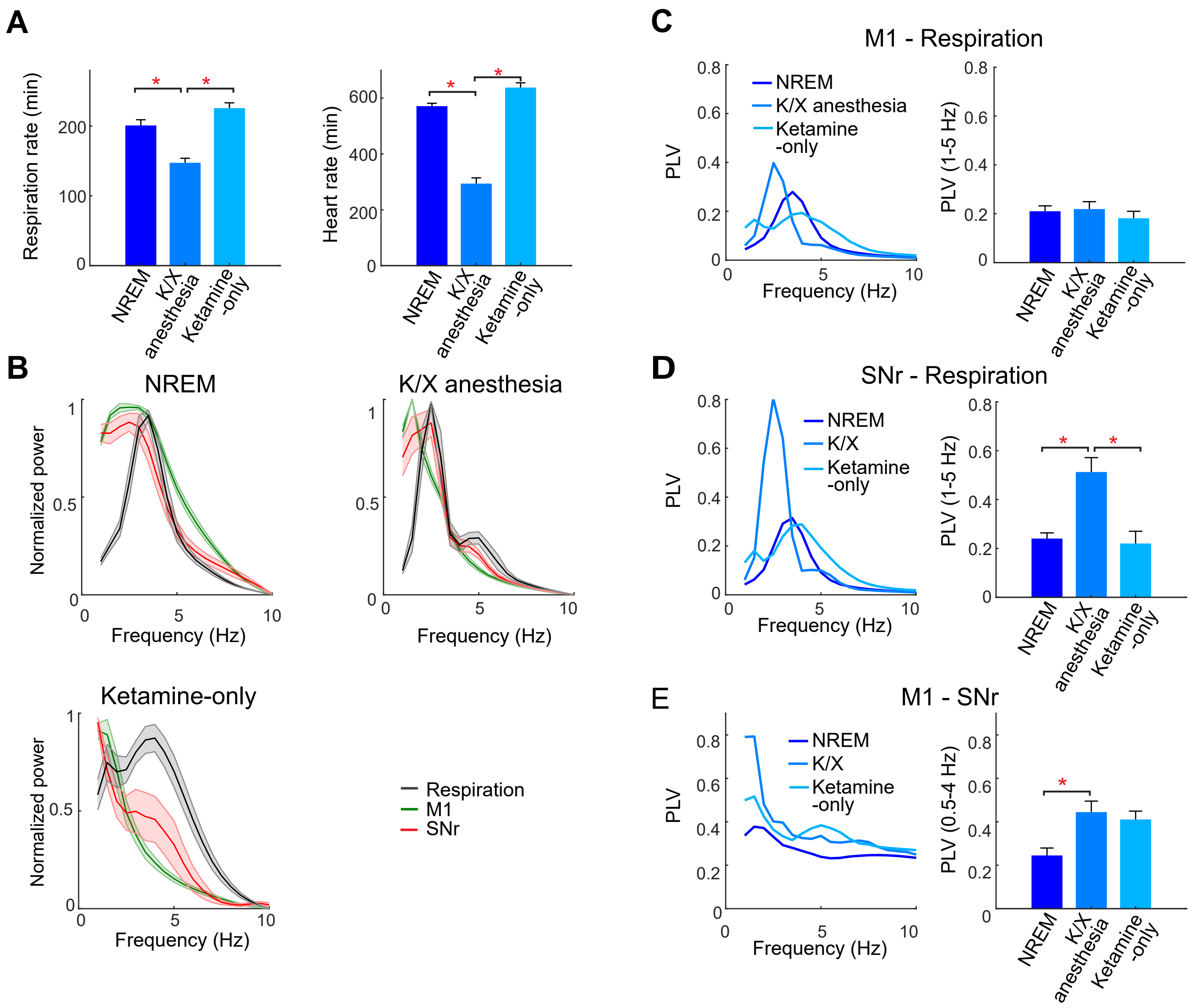
